# Delineating and validating higher-order dimensions of psychopathology in the Adolescent Brain Cognitive Development (ABCD) study

**DOI:** 10.1101/548057

**Authors:** Giorgia Michelini, Deanna M. Barch, Yuan Tian, David Watson, Daniel N. Klein, Roman Kotov

## Abstract

Hierarchical dimensional systems of psychopathology promise more informative descriptions for understanding risk and predicting outcome than traditional diagnostic systems, but it is unclear how many major dimensions they should include. We delineated the hierarchy of childhood and adult psychopathology and validated it against clinically-relevant measures. Participants were 4,524 9- and 10-year-old children and their parents from the Adolescent Brain Cognitive Development (ABCD) study. Factor analyses on items from the Child Behavior Checklist and Adult Self-Report characterized a dimensional hierarchy. We examined the familial aggregation of the psychopathology dimensions, and the ability of different factor solutions to account for risks factors, social, educational and cognitive functioning, and physical and mental health service utilization. A hierarchical structure with a general psychopathology (‘p’) factor at the apex and five specific factors (internalizing, somatoform, detachment, neurodevelopmental, and externalizing) emerged in children. Adult factors were similar, but externalizing behaviors separated into disinhibited and antagonistic factors. Child and parent p-factors correlated highly (*r*=.61, *P*<.001), and smaller but significant correlations emerged for convergent dimensions between parents and children after controlling for p-factors (*r*=.10-20, *P*<.001). A model with childhood p-factor alone explained mental health service utilization (R^2^=.13, *P*<.001), but up to five dimensions provided incremental validity to account for developmental risk and current functioning (R^2^=.03-.20, *P*<.001). In this first investigation comprehensively mapping the psychopathology hierarchy in children and adults, we delineated a hierarchy of higher-order dimensions associated with a range of clinically-relevant validators. These findings hold important implications for psychiatric nosology and future research in this sample.

## 1. INTRODUCTION

Traditional psychiatric nosologies define mental disorders as distinct categories,^1, 2^ but this is at odds with extensive evidence that disorders lie on a continuum with normality and are highly comorbid.^3–7^ This comorbidity reflects underlying higher-order dimensions (or spectra) of psychopathology.^4, 7–9^ Dimensional classifications of these spectra have been proposed as alternative approaches to better align the nosology with empirical evidence.^4, 7, 8, 10^ However, available models differ in the number of spectra that they specify.

Numerous studies point to a general factor (‘p’) that represents common susceptibility to psychopathology and explains why all mental disorders tend to co-occur.^5, 9, 11–14^ Other research supports a separation between broad internalizing and externalizing spectra – originally identified in studies that shaped the Achenbach System of Empirically Based Assessment (ASEBA)^15, 16^ – arguing that this is an important distinction both in adults^17, 18^ and children.^19, 20^ However, further evidence suggests that a greater number of major dimensions are needed to characterize psychopathology.^8, 21–25^ For instance, the recently-developed Hierarchical Taxonomy of Psychopathology (HiTOP)^7, 8^ includes six spectra (internalizing, somatoform, detachment, thought disorder, antagonism, and disinhibition), which were identified based on extensive factor analytic literature (for a review see^8^). Yet, the dimensions depicted in these studies may not provide full coverage of psychopathology. For example, a neurodevelopmental spectrum (e.g., speech problems, motor problems, autism) has been proposed,^26^ but is still undergoing factor analytic examination on its placement among other forms of psychopathology.^27, 28^

Models with different numbers of dimensions remain to be reconciled in order to advance psychiatric classification and its clinical utility. Simpler and more complex architectures may be integrated as different levels of a single hierarchy: from a p-factor at the apex to progressively more specific nested factors.^29, 30^ Consequently, models with different numbers of dimensions (one, two, three etc.) can co-exist and be studied simultaneously. Initial studies have identified a hierarchy of higher-order dimensions,^22, 23, 30–32^ but were largely limited to personality pathology and focused on adults. Importantly, developmental studies suggest that there may be progressive differentiation in psychopathology and emergence of additional dimensions over development^14, 33^ which underscores the importance of studying children samples as well.

Beyond the identification of the number of dimensions, an important step for delineating a new psychopathology classification is to validate dimensions against criteria important for clinical practice and research, such as genetic/familial and psychosocial risk factors, cognitive processes, illness course, and treatment outcome.^34–36^ In a hierarchical structure, validity may differ across levels, as more elaborate models tend to be more informative, but are less parsimonious, and the choice between models may depend on the purpose of inquiry. Available studies show that broader spectra are associated with familiality for psychiatric disorders, childhood adversities, brain and functional impairment^11, 13, 35^ while more specific dimensions are required to adequately account for outcomes such as educational achievement and executive functioning.^14, 19, 27^ However, a systematic evaluation of validity of dimensions across hierarchical levels is lacking.

In the present study, we sought to delineate higher-order dimensions of psychopathology within a hierarchical structure, and compare the validity of different levels of specificity. Our first aim was to investigate the hierarchical structure of psychopathology in 4,524 children from the Adolescent Brain Cognitive Development (ABCD) study^37–39^ – and attempt to replicate it in their parents – by analyzing a large and diverse set of symptoms.^15, 16^ Our second aim was to compare the validity of different levels of the childhood psychopathology hierarchy in relation to clinically-informative measures of familial and developmental risk factors, current social, academic and cognitive functioning, and service utilization.^11, 14, 36, 40, 41^

## 2. METHODS

### 2.1. Sample

The sample for this study consisted of children enrolled in the ABCD study and their parents. The ABCD study is a collaboration between 21 sites across the US to investigate psychological and neurobiological development from preadolescence to early adulthood. Full details on recruitment can be found elsewhere.^37^ Briefly, the primary method for recruiting children aged 9 or 10 at the time of the baseline assessments (between 2016 and 2018) and their parents was probability sampling of public and private elementary schools within the catchment areas of the 21 research sites, encompassing over 20% of the entire US population of 9-10 year olds. School selection was based on gender, race and ethnicity, socioeconomic status, and urbanicity. No exclusion criteria were applied other than age and attending a public or private elementary school in the catchment area, with the aim to recruit a cohort representative of the sociodemographic variation in the US population. The cohort’s representation of diverse demographic and socio-economic groups was monitored through the National Center for Education Statistics databases, containing socio-demographic characteristics of the students attending each school. This enabled dynamic adjustment of the accumulating sample based on demographic targets throughout recruitment. The present study is based on 4,524 children (47.57% females) and 4,524 parents (one per child; mean age=40.5, SD=6.69; 85.06% females) from the first data release (NDAR-DOI: 10.15154/1412097). Demographic characteristics for participants in this data release have been reported elsewhere.^42^ All procedures were approved at each site by institutional ethics boards. Parents or guardians provided written informed consent after the procedures had been fully explained and children assented before participation in the study.^43^

### 2.2. Measures

Full details on measures are presented in Supplementary Method 1. Children and parents completed assessments during an in-person visit. Psychopathology was examined in the children with the parent-reported Child Behavior Checklist (CBCL)^15^ and in the adults with the Adult Self-Report (ASR)^16^ from ASEBA, which assess problems occurring in the past 6 months on a 3-point scale.

For validation, we examined key risk factors, functioning measures, and service use variables: history of developmental motor and speech delays,^38^ conflict within the family, social peer functioning (number of friends), academic functioning (school connectedness, average grades),^44^ cognitive functioning (crystalized and fluid intelligence composites from the National Institute of Health Toolbox),^39^ utilization of physical and mental health services, and current use of medications.^38^

### 2.3. Statistical analysis

To investigate the hierarchical structure of psychopathology, we employed an exploratory approach, given uncertainties regarding the number of dimensions and the composition of the levels of the hierarchy. Specifically, we used exploratory factor analysis (EFA) to empirically extract (with principal component analysis) and rotate (with promax) factor solutions with an increasing number of factors. To avoid distorting the factor structure in EFA with items that were not analyzable due to being endorsed too infrequently or too-highly correlated with other items,^45^ we removed items which frequency was too low (<1% rated as 1 or 2) and aggregated items that were highly correlated (polychoric r>.75) into composites (see Supplementary Method 1). The maximum number of factors to extract was determined with parallel analyses^46^ (extraction was stopped when eigenvalues fell within the 95% confidence interval of eigenvalues from simulated data; Supplementary Figure 1). Since parallel analysis has a tendency to over-factor, we also examined the interpretability of factor solutions,^45, 47^ defined as presence of >3 clear primary loadings (highest loading ≥.35 and at least .10 greater than all other loadings) for each factor.^45, 47^ All factor structures from one to the maximum number of factors were considered. To map the hierarchical structure, we correlated factor scores on adjacent levels of the hierarchy to describe transitions between levels using Goldberg’s hierarchical method,^29^ in line with previous work.^30, 48, 49^ The paths between levels in the hierarchical model reflect correlations ≥.55 between the factor scores where a ≥ shift of at least two primary loadings was observed between levels.

To compare utility of the solutions, in validation analyses we first examined the degree of familial aggregation of the dimensions by correlating the factor scores derived for each dimensions in parents and children, using both zero-order correlations and partial correlations controlling for the first general psychopathology factors in parents and children. To familial aggregation due to shared genetic and environmental factors between parent and child, 194 non-biological parent-child pairs were excluded from this analysis (112 adoptive parents, 30 custodial parents, 52 other non-biological parents). Second, we entered the factor scores from each level of the childhood hierarchy as separate blocks into a hierarchical regression model, with each of the validators as the independent variable. We examined the predictive power and the incremental validity of each level of the hierarchy over more parsimonious structures with the significance of R^2^ change between blocks.^48^ We used this stringent test, rather than comparing levels in pairs, to ensure that a significant result for models with more factors reflects new information not captured by simpler factor solutions. All analyses were run in Mplus version 7 (Muthén and Muthén, Los Angeles, CA) and SPSS version 25 (IBM Corp, Armonk, NY).

## 3. RESULTS

### 3.1. Hierarchical factor structure of CBCL and ASR

#### 3.1.1. CBCL

Parallel analyses indicated that up to 17 factors could be extracted from CBCL items (Supplementary Figure 1). After examining interpretability of these factor solutions, 1- to 5-factor solutions were found to be acceptable (Table 1, Supplementary Table 1). Solution with more than five factors were not tenable as each included at least one factor without any primary loadings (e.g., Supplementary Table 1).

**Table 1.** Factor loadings (top) and factor correlations (bottom) for the 5-factor solution from the exploratory factor analysis on CBCL items.

| CBCL items | F1 | F2 | F3 | F4 | F5 |
| --- | --- | --- | --- | --- | --- |
| **<u>Primary-loading items</u>** |  |  |  |  |  |
| Composite (Attacks/Threatens) | **.86** | -.08 | .03 | -.05 | .04 |
| Cruelty, bullying, or meanness to others | **.85** | -.07 | -.06 | -.04 | -.02 |
| Composite (Disobeys rules) | **.78** | .22 | -.08 | -.04 | -.05 |
| Gets in many fights | **.76** | .10 | -.10 | .05 | -.02 |
| Temper tantrums or hot temper | **.75** | -.05 | .24 | -.01 | -.09 |
| Argues a lot | **.73** | .10 | .13 | -.02 | -.14 |
| Doesn't seem to feel guilty after misbehaving | **.71** | .11 | -.12 | -.09 | .08 |
| Composite (Steals) | **.71** | .06 | -.20 | .04 | .13 |
| Screams a lot | **.70** | -.01 | .17 | .01 | -.04 |
| Swearing or obscene language | **.67** | -.03 | -.04 | .01 | .08 |
| Runs away from home | **.69** | -.08 | .09 | .12 | .11 |
| Lying or cheating | **.67** | .13 | -.16 | .08 | .02 |
| Stubborn, sullen, or irritable | **.67** | -.05 | .24 | .07 | .00 |
| Composite (Destroys) | **.67** | .18 | -.04 | .03 | .06 |
| Cruel to animals | **.67** | -.01 | -.04 | -.04 | .08 |
| Teases a lot | **.62** | .12 | -.03 | .07 | -.12 |
| Easily jealous | **.61** | -.01 | .30 | -.07 | -.05 |
| Sudden changes in mood or feelings | **.58** | -.05 | .32 | .05 | .10 |
| Suspicious | **.56** | .02 | .25 | -.01 | .15 |
| Sulks a lot | **.54** | -.08 | .30 | .07 | .15 |
| Bragging, boasting | **.51** | .18 | .02 | .09 | -.27 |
| Thinks about sex too much | **.50** | .18 | .09 | -.02 | -.09 |
| Showing off or clowning | **.50** | .33 | -.07 | .05 | -.27 |
| Hangs around with others who get in trouble | **.49** | .20 | -.12 | -.02 | .01 |
| Demands a lot of attention | **.48** | .28 | .27 | -.01 | -.22 |
| Composite (Peer problems) | **.47** | .20 | .04 | .00 | .26 |
| Whining | **.40** | .09 | .24 | .13 | -.07 |
| Composite (Distracted) | .11 | **.86** | -.06 | -.05 | .05 |
| Can't sit still, restless, or hyperactive | .23 | **.75** | .04 | -.10 | -.18 |
| Daydreams or gets lost in his/her thoughts | -.16 | **.61** | .03 | .10 | .21 |
| Stares blankly | -.08 | **.61** | -.03 | .10 | .36 |
| Confused or seems to be in a fog | -.05 | **.58** | .10 | .04 | .35 |
| Nervous movements or twitching | -.10 | **.57** | .35 | -.06 | -.02 |
| Poorly coordinated or clumsy | -.07 | **.55** | .00 | .20 | .17 |
| Poor school work | .25 | **.55** | -.15 | -.05 | .21 |
| Fails to finish things he/she starts | .25 | **.54** | -.03 | .01 | .11 |
| Talks too much | .21 | **.51** | .01 | .14 | -.24 |
| Repeats certain acts over and over; compulsions | .13 | **.50** | .17 | -.05 | .07 |
| Can't get his/her mind off certain thoughts; obsessions | .12 | **.48** | .30 | -.01 | .01 |
| Acts too young for his/her age | .17 | **.47** | .05 | -.08 | .13 |
| Strange behavior | .25 | **.43** | .12 | .01 | .24 |
| Strange ideas | .22 | **.38** | .03 | .04 | .23 |
| Too fearful or anxious | -.13 | .27 | **.78** | .01 | -.01 |
| Worries | -.03 | .06 | **.67** | .19 | .00 |
| Feels too guilty | -.01 | .15 | **.64** | .12 | .00 |
| Feels he/she has to be perfect | .01 | -.10 | **.63** | .07 | .00 |
| Nervous, high-strung, or tense | .02 | .37 | **.62** | -.01 | -.06 |
| Fears he/she might think or do something bad | .08 | .10 | **.55** | .06 | .02 |
| Feels worthless or inferior | .21 | .06 | **.53** | .00 | .20 |
| Self-conscious or easily embarrassed | .09 | -.02 | **.47** | .07 | .29 |
| Fears certain animals, situations, or places, other than school | -.05 | .20 | **.46** | .01 | .08 |
| Fears going to school | .06 | .08 | **.44** | .13 | .19 |
| Clings to adults or too dependent | .07 | .25 | **.37** | .00 | .12 |
| Nausea, feels sick | -.01 | -.07 | .07 | **.88** | -.02 |
| Stomachaches | .00 | -.07 | .08 | **.79** | -.05 |
| Vomiting, throwing up | -.01 | -.04 | -.20 | **.75** | .08 |
| Headaches | .03 | -.01 | .00 | **.63** | -.01 |
| Aches or pains (not stomach or headaches) | .03 | .05 | .05 | **.53** | .01 |
| Feels dizzy or lightheaded | -.05 | .09 | .20 | **.53** | .06 |
| Rashes or other skin problems | .03 | .07 | .07 | **.36** | .01 |
| Withdrawn, doesn't get involved with others | .20 | .01 | .12 | -.02 | **.71** |
| Would rather be alone than with others | .07 | .09 | .11 | .02 | **.58** |
| Too shy or timid | -.07 | -.05 | .36 | -.04 | **.53** |
| Refuses to talk | .25 | .04 | .14 | -.03 | **.48** |
| Underactive, slow moving, or lacks energy | .04 | .17 | -.04 | .30 | **.48** |
| **<u>Non-primary loading or cross-loading items</u>** |  |  |  |  |  |
| Cries a lot | .28 | .13 | .31 | .10 | .04 |
| Feels or complains that no one loves him/her | .49 | -.05 | .44 | -.07 | .12 |
| Talks about killing self | .39 | .02 | .37 | .03 | .15 |
| Unusually loud | .36 | .42 | .08 | .12 | -.19 |
| Impulsive or acts without thinking | .46 | .51 | .03 | -.03 | -.10 |
| Prefers being with older kids | .31 | .17 | -.06 | .11 | .07 |
| Complains of loneliness | .25 | .15 | .32 | .02 | .22 |
| Feels others are out to get him/her | .42 | .06 | .39 | -.04 | .13 |
| Gets teased a lot | .22 | .26 | .08 | .09 | .29 |
| Gets hurt a lot, accident prone | .08 | .31 | -.02 | .33 | -.03 |
| Prefers being with younger kids | .11 | .33 | .06 | .03 | .19 |
| Speech problem | -.02 | .34 | -.06 | .04 | .22 |
| Constipated, doesn't move bowels | .03 | .12 | .15 | .25 | .13 |
| Overtired without good reason | .09 | .13 | .11 | .34 | .30 |
| Nightmares | .07 | .22 | .24 | .29 | -.04 |
| Deliberately harms self or attempts suicide | .35 | -.02 | .41 | .03 | .17 |
| Composite (Hallucinations) | .13 | .15 | .25 | .11 | .11 |
| Picks nose, skin, or other parts of body | .16 | .31 | .04 | .11 | -.02 |
| Composite (Play sex) | .27 | .20 | .06 | -.04 | .02 |
| Sleeps less than most kids | .12 | .27 | .13 | .13 | .14 |
| Stores up too many things he/she doesn't need | .18 | .20 | .14 | .07 | .09 |
| Talks or walks in sleep | .02 | .20 | .06 | .28 | -.17 |
| Trouble sleeping | .08 | .28 | .25 | .23 | .02 |
| Secretive, keeps things to self | .34 | .02 | .10 | .03 | .38 |
| There is very little he/she enjoys | .40 | .08 | .13 | -.03 | .36 |
| Unhappy, sad, or depressed | .35 | -.06 | .39 | .12 | .27 |
| Bowel movements outside toilet | .12 | .11 | -.11 | .23 | .28 |
| Bites fingernails | .06 | .25 | .08 | .06 | -.02 |
| Doesn't eat well | .19 | .18 | .06 | .07 | .15 |
| Sleeps more than most kids during day and/or night | .02 | .13 | .09 | .14 | .20 |
| Wets the bed | .05 | .20 | -.13 | .12 | .08 |
| Wets self during the day | -.02 | .33 | -.05 | .20 | .19 |
| Composite (Weight problems) | .11 | .04 | -.05 | .19 | .21 |
| Thumb-sucking | .21 | -.07 | .01 | .04 | -.04 |

**Factor correlations**
|  | **F1** | **F2** | **F3** | **F4** | **F5** |
| --- | --- | --- | --- | --- | --- |
| F1 (Externalizing) | 1 |  |  |  |  |
| F2 (Neurodevelopmental) | .55 | 1 |  |  |  |
| F3 (Internalizing) | .39 | .35 | 1 |  |  |
| F4 (Somatoform) | .34 | .34 | .41 | 1 |  |
| F5 (Detachment) | .32 | .30 | .35 | .27 | 1 |

Notes: Bold indicates primary loadings ( $\geq .35$ ) with at least .10 difference from the second largest loading.
All factor correlations were statistically significant ( $P < .001$ ). CBCL=Child Behavior Checklist,
F1=Externalizing factor, F2=Neurodevelopmental factor, F3=Internalizing factor, F4=Somatoform factor,
F5=Detachment factor.

All models from 1-factor to 5-factor were interpretable and are represented as a hierarchical structure (Figure 1), with paths showing correlations between levels. The 1-factor structure reflected a general childhood psychopathology p-factor.^5, 14^ The 2-factor solution revealed the expected broad internalizing and broad externalizing factors.^15, 19, 50^ In the 3-factor structure, the broad externalizing factor split into externalizing (e.g. rule-breaking and aggressive behavior) and neurodevelopmental factors (e.g. inattention, hyperactivity, impulsivity, clumsiness, speech problems). In the 4-factor solution, somatoform problems emerged from the broad internalizing factor. In the 5-factor structure, the broad internalizing factor split into narrower internalizing problems (e.g. anxiety, depressive symptoms) and detachment. Factors in the final 5-factor solution showed small-to-large correlations with one another (*r*=.27-.55) (Table 1).

**Figure 1.**
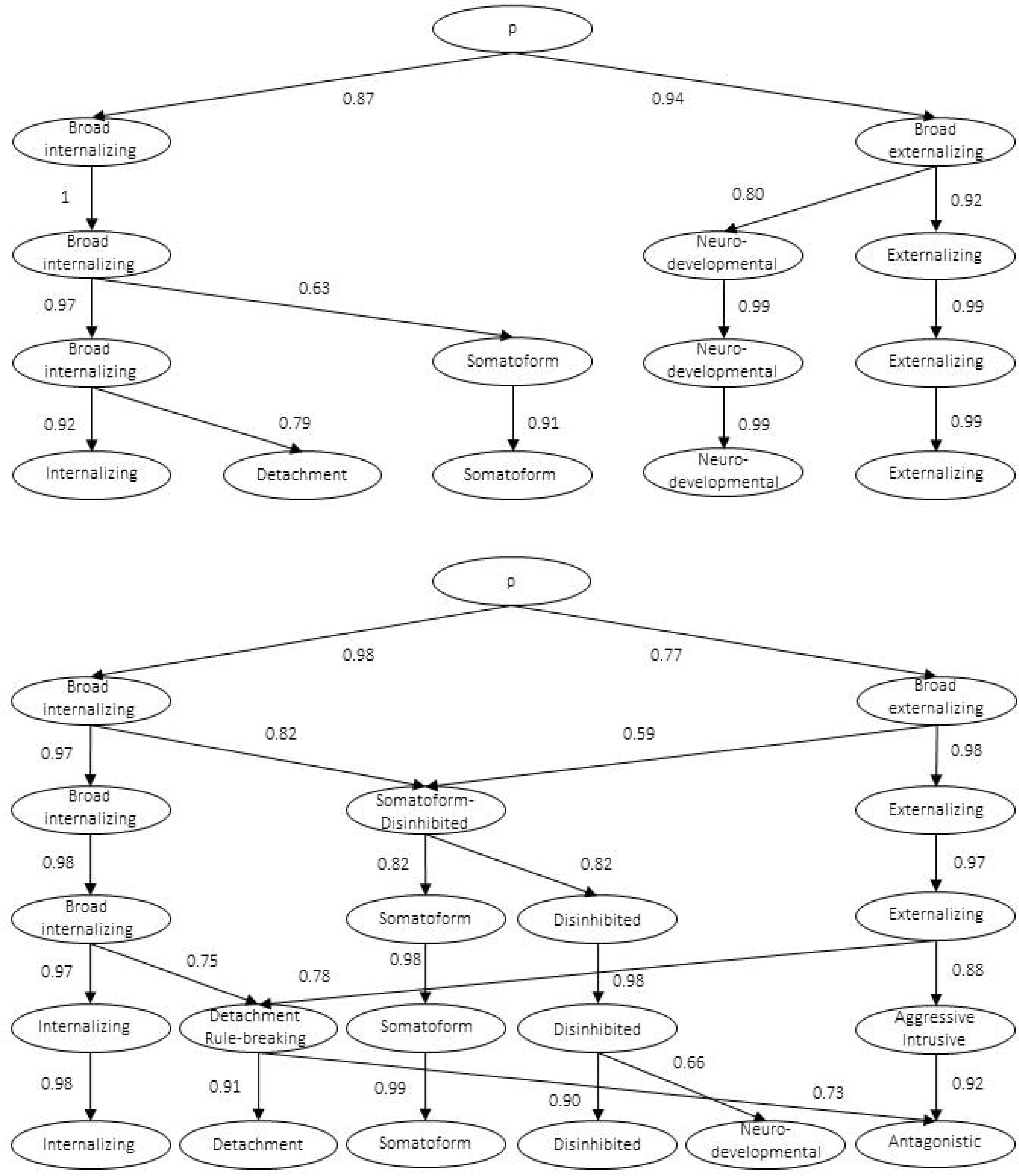
Hierarchical models from CBCL items (top half) and ASR items (bottom half) illustrating hierarchies of child and adult psychopathology. Note: CBCL=Childhood Behavior Checklist, ASR=Adult Self Report.

#### 3.1.2. ASR

Parallel analyses indicated that up to 17 factors could be extracted from ASR items (Supplementary Figure 1). The 6-factor solution was the most differentiated interpretable structure (Table 2, Supplementary Table 2), as factor solutions with more factors were not interpretable. For example, the last factor in the 7- and 8-factor models included only two-to-three primary loadings, thus indicating no other meaningful factors beyond six (Supplementary Table 2).

**Table 2.**
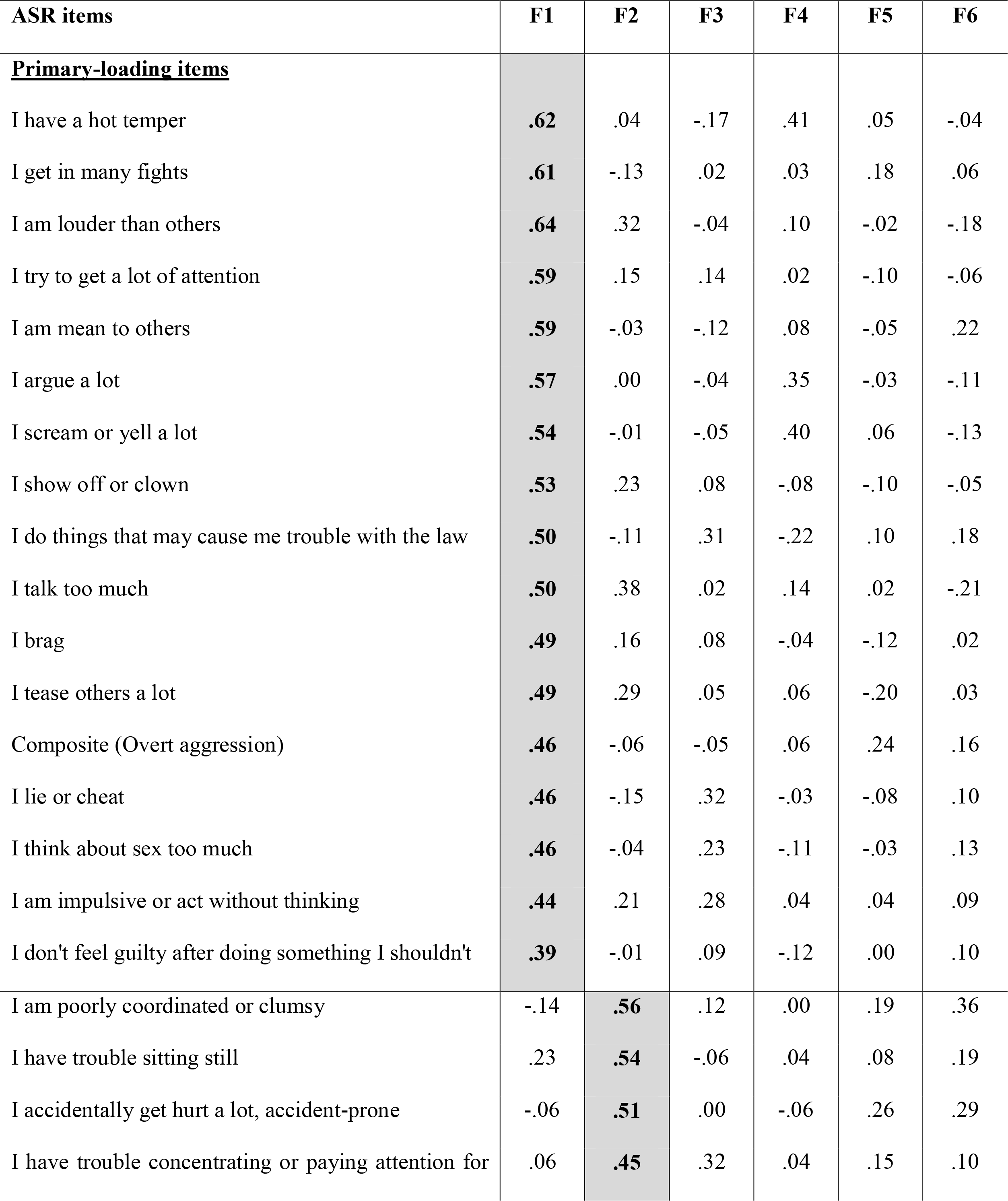
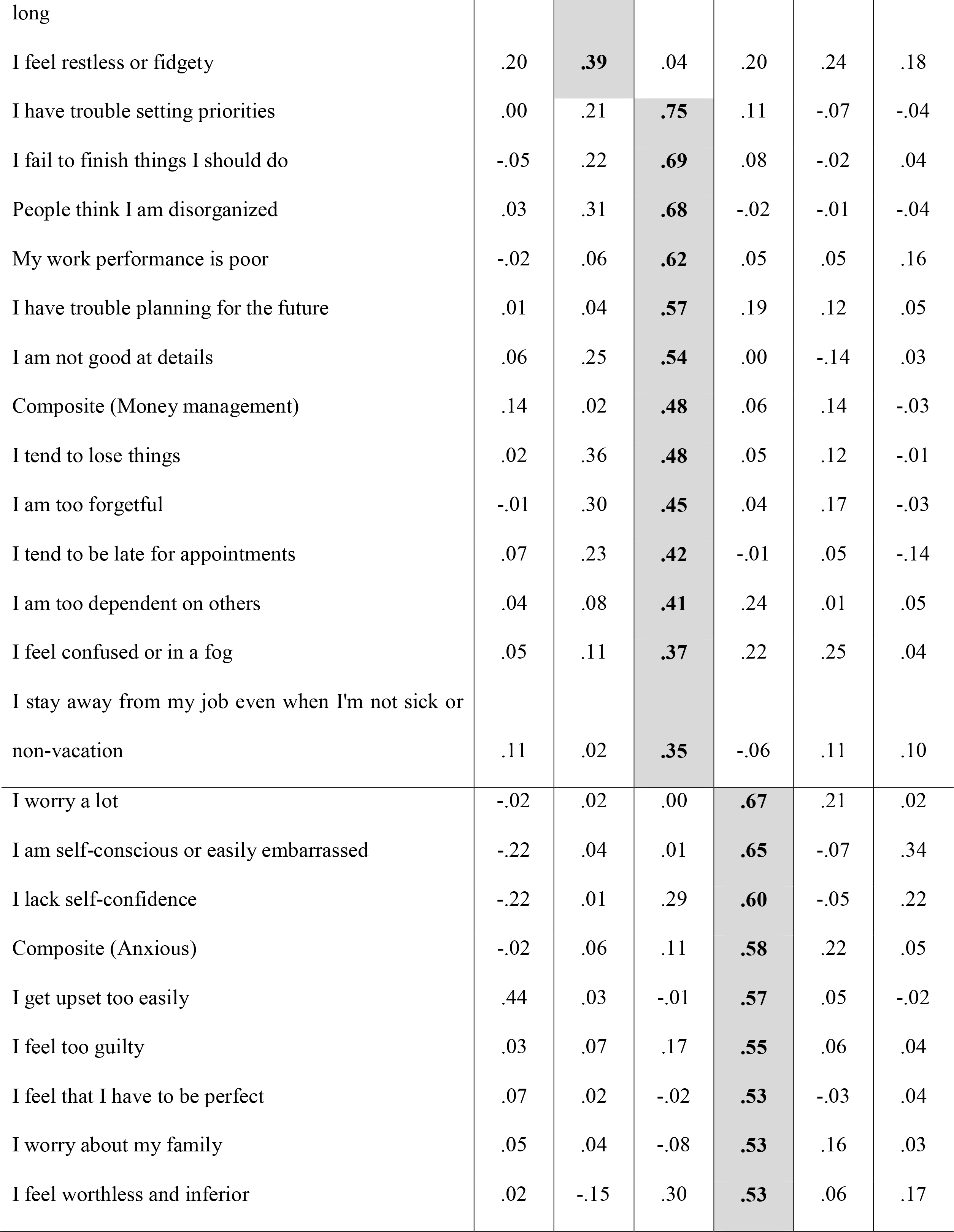
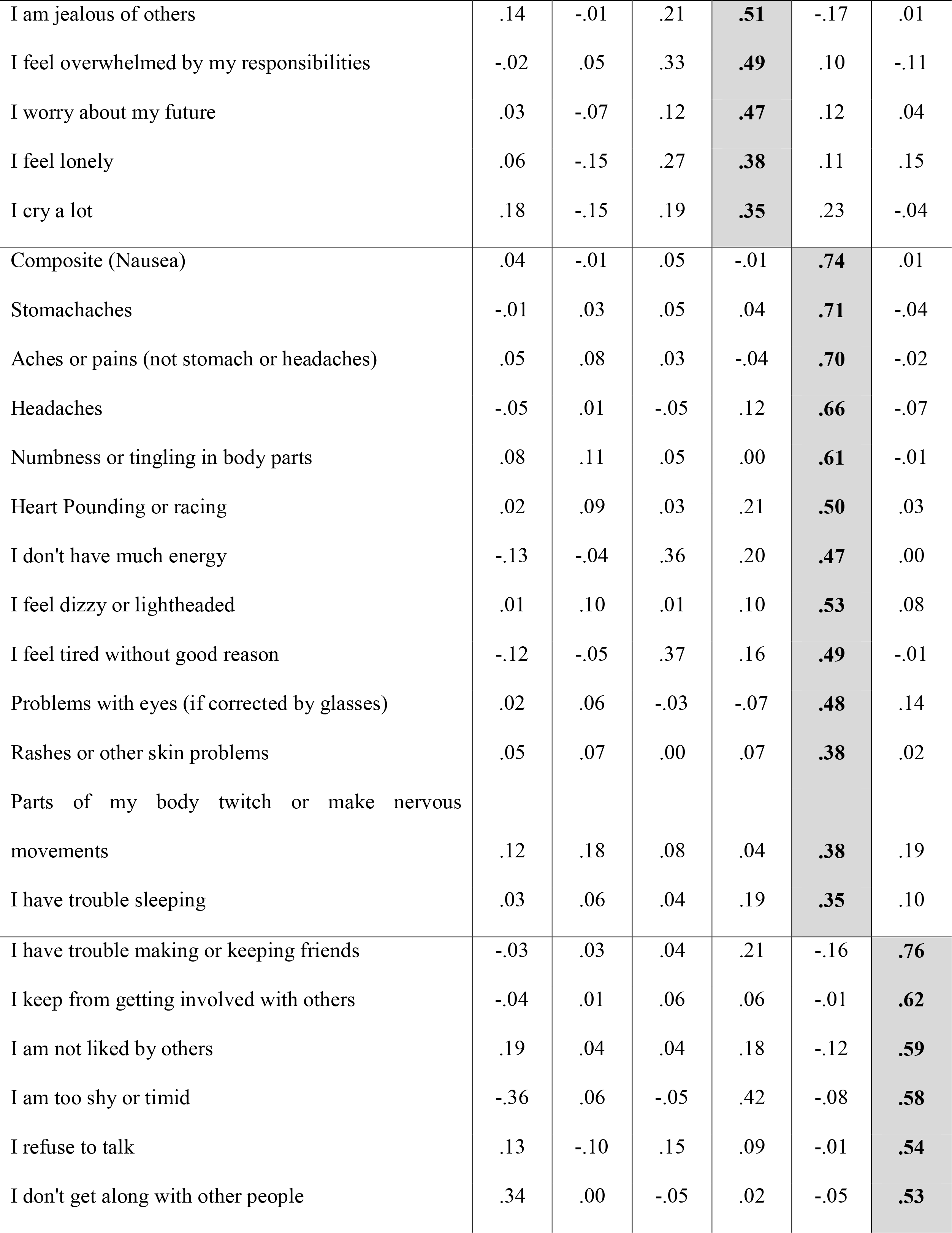
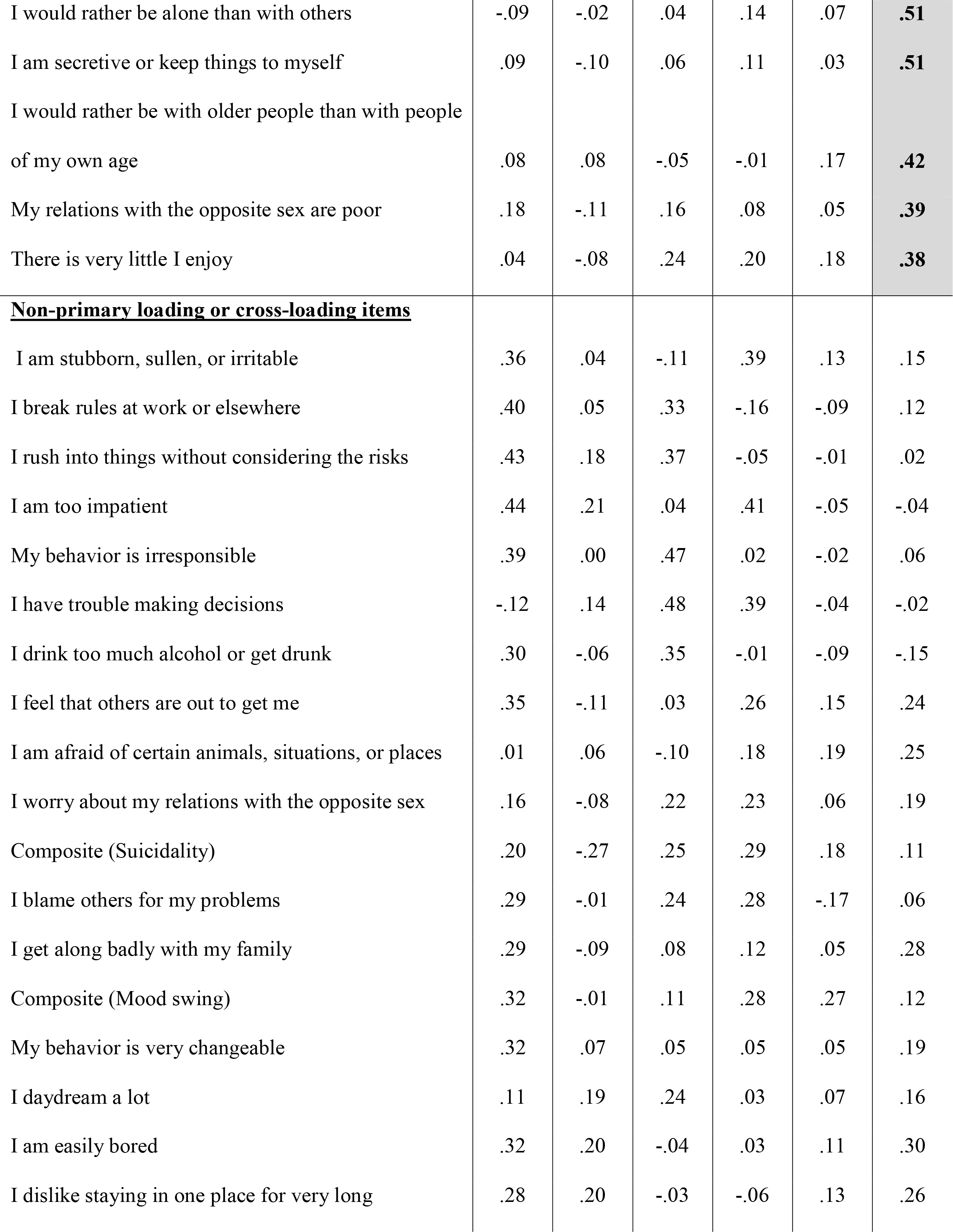
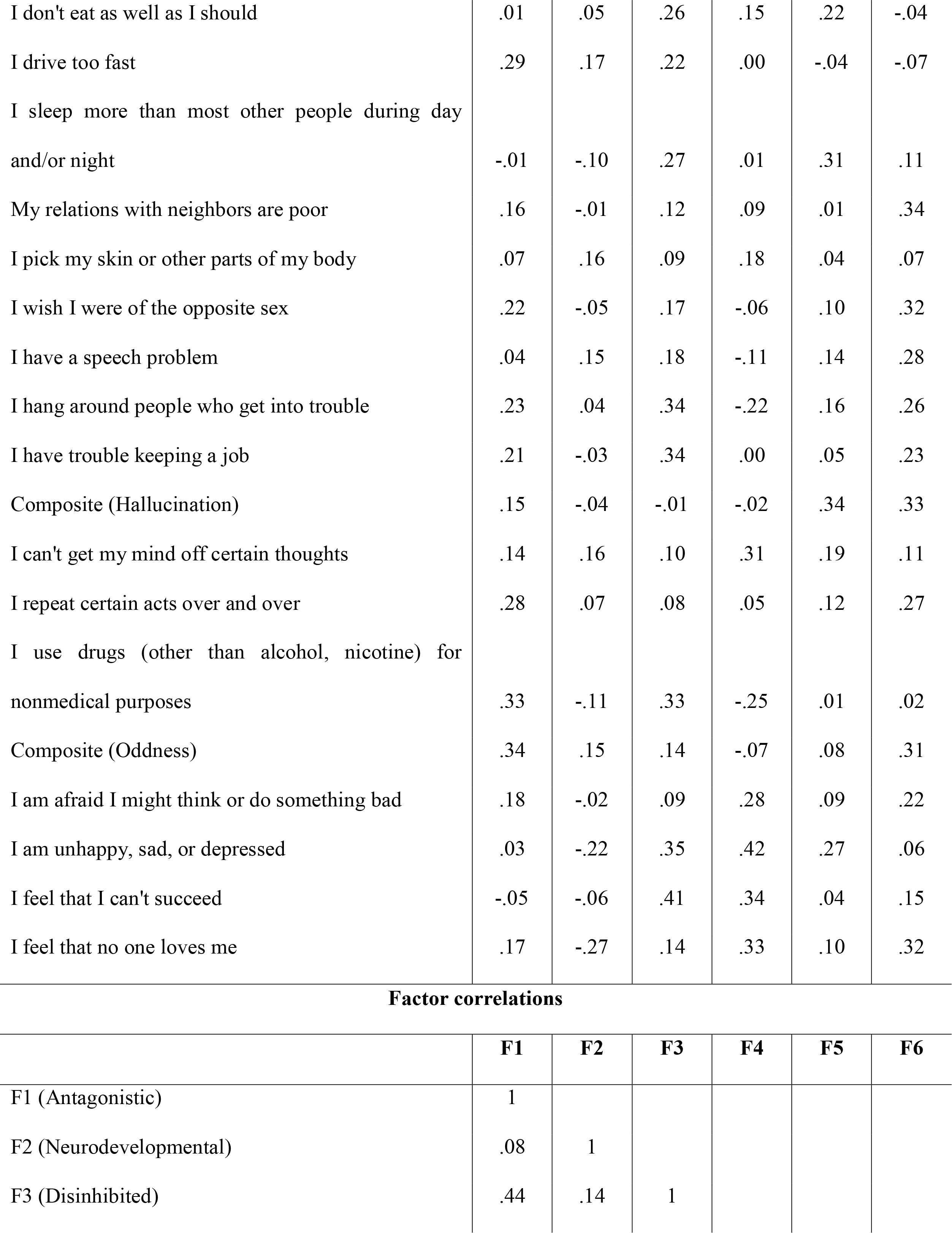
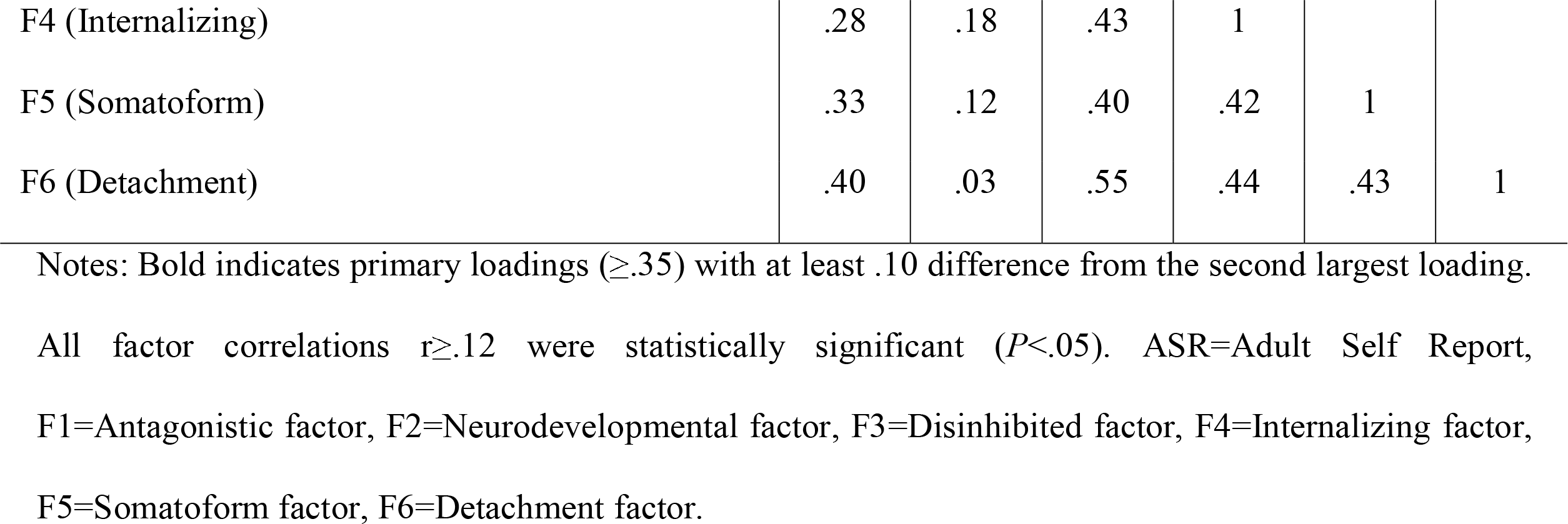
Factor loading (top) and factor correlations (bottom) for the 6-factor solution from the exploratory factor analysis on ASR items.

All models from 1-factor to 6-factor are represented in Figure 1. The 1-factor structure reflected p-factor.^11^ The 2-factor solution showed the broad internalizing and externalizing factors.^17^ In the 3-factor structure, a factor encompassing disinhibited (e.g. poor planning) and somatoform problems (e.g. complains about aches) originated from the broad internalizing and externalizing factors. In the 4-factor solution, the somatoform-disinhibited factor split into separate disinhibited and somatoform dimensions. In the 5-factor structure, externalizing and broad internalizing factors reorganized into detachment/rule-breaking, aggressive-intrusive, and internalizing factors. In the 6-factor solution, the disinhibited factor split into disinhibited and neurodevelopmental (e.g. clumsiness, hyperactivity) dimensions; and rule-breaking behaviors joined aggressive and intrusive behaviors to form an antagonistic factor, leaving a distinct detachment dimension. Factors in the final 6-factor solution showed small-to-large correlations with one another (*r*=.08-.55) (Table 2).

### 3.2. Validation analyses

#### 3.2.1. Familial aggregation

Zero-order correlations between the five childhood and six adult factor scores ranged between *r*=.14-.44, *P*<.001 (Table 3). The correlation between child and parent p-factor scores was *r*=.61, *P*<.001. This pattern suggested substantial familial aggregation of a common vulnerability, explaining co-occurrence across psychopathology dimensions. Controlling for these two p-factors revealed a more specific pattern of familial aggregation between corresponding parent and child dimensions, except for child externalizing with parent disinhibition (largely encompassing adult symptoms not included in the childhood measure). All other convergent partial correlations ranged between *r*=.10-.19 (*P*<.001) and were significantly larger than all discriminant partial correlations, based on Fisher’s z tests (Table 3).

**Table 3.**
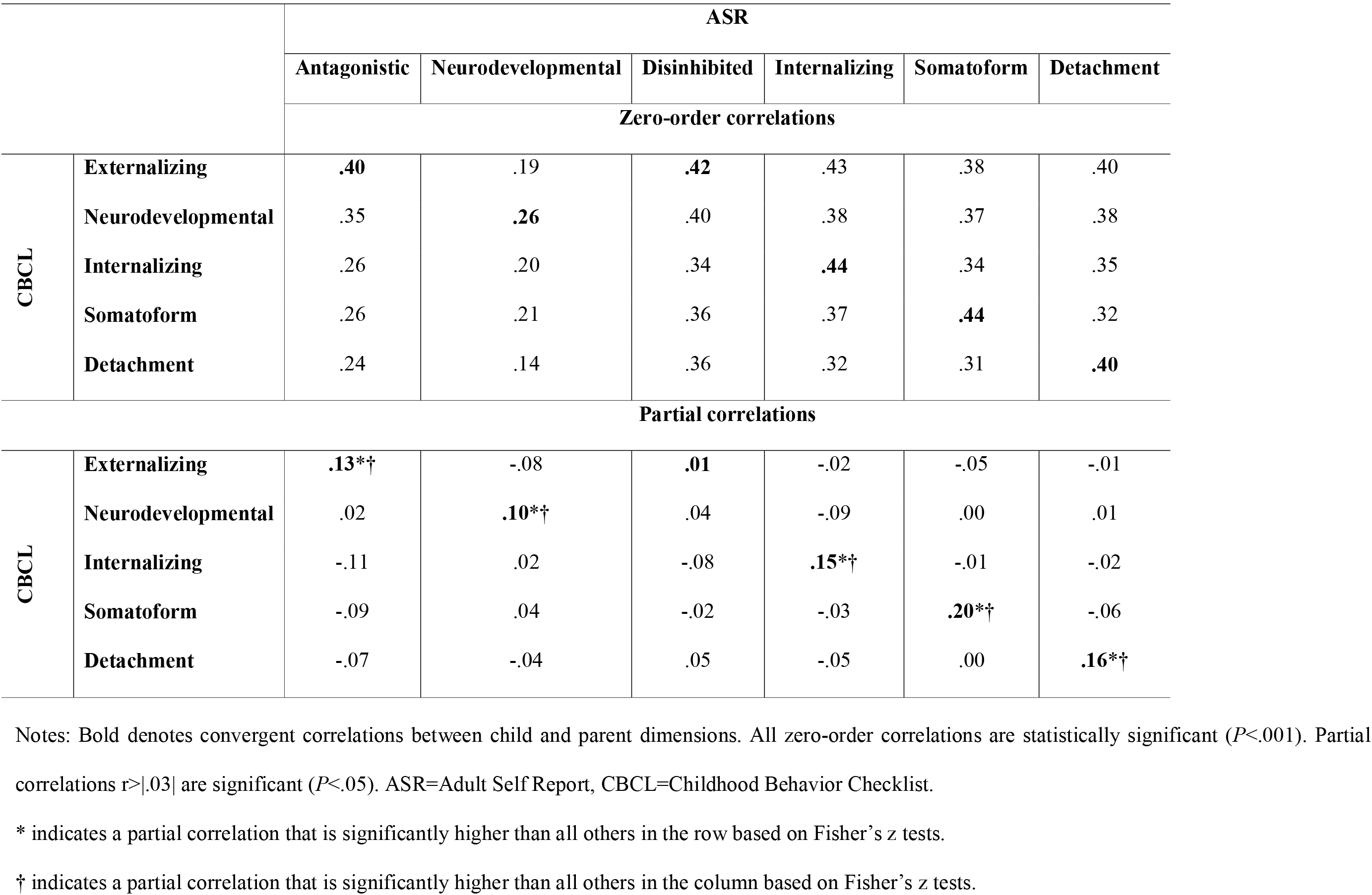
Zero-order (top half) and partial correlations (bottom half) between the dimensions in the 5- and 6-factor structures from CBCL and ASR items, respectively, controlling for childhood and adult p-factors.

|  |  | ASR |  |  |  |  |  |
| --- | --- | --- | --- | --- | --- | --- | --- |
|  |  | Antagonistic | Neurodevelopmental | Disinhibited | Internalizing | Somatoform | Detachment |
|  |  | Zero-order correlations |  |  |  |  |  |
| CBCL | Externalizing | **.40** | .19 | **.42** | .43 | .38 | .40 |
|  | Neurodevelopmental | .35 | **.26** | .40 | .38 | .37 | .38 |
|  | Internalizing | .26 | .20 | .34 | **.44** | .34 | .35 |
|  | Somatoform | .26 | .21 | .36 | .37 | **.44** | .32 |
|  | Detachment | .24 | .14 | .36 | .32 | .31 | **.40** |
| Partial correlations |  |  |  |  |  |  |  |
| CBCL | Externalizing | **.13\*†** | -.08 | **.01** | -.02 | -.05 | -.01 |
|  | Neurodevelopmental | .02 | **.10\*†** | .04 | -.09 | .00 | .01 |
|  | Internalizing | -.11 | .02 | -.08 | **.15\*†** | -.01 | -.02 |
|  | Somatoform | -.09 | .04 | -.02 | -.03 | **.20\*†** | -.06 |
|  | Detachment | -.07 | -.04 | .05 | -.05 | .00 | **.16\*†** |

Notes: Bold denotes convergent correlations between child and parent dimensions. All zero-order correlations are statistically significant ( $P < .001$ ). Partial correlations $r > .03$ are significant ( $P < .05$ ). ASR=Adult Self Report, CBCL=Childhood Behavior Checklist. \* indicates a partial correlation that is significantly higher than all others in the row based on Fisher's z tests. † indicates a partial correlation that is significantly higher than all others in the column based on Fisher's z tests.

#### 3.2.2. Validity of childhood hierarchical structure

The 1-factor solution was significantly associated with all validators (Figure 2, Supplementary Table 3). The p-factor alone explained 13.43% of the variance in utilization of mental health services, and addition of more differentiated factors, although statistically significant, produced minimal improvement in R^2^ (up to 14.38%). For medication use, medical history, family conflict, and school connectedness, the p-factor alone explained 2.57-2.95% of the variance, and the addition of more complex factor structures provided a moderate increase, contributing up to 4.02-4.93% of variance. For crystalized intelligence and fluid intelligence, the 1-factor model predicted only a small proportion of variance (.60% and 1.79% respectively), with a substantial increase in R^2^ by adding the 2-, 3- and 4-factor solutions (up to 3.46 and 4.18% respectively), but not the 5-factor solution. The 1-factor accounted for a relatively small proportion of the variance compared to the more complex factor solutions for average grades (from 7.81% for p-factor to 19.34% total), number of friends (.12% to 3.28%), and history of developmental delays (.37% to 2.94%).

**Figure 2.**
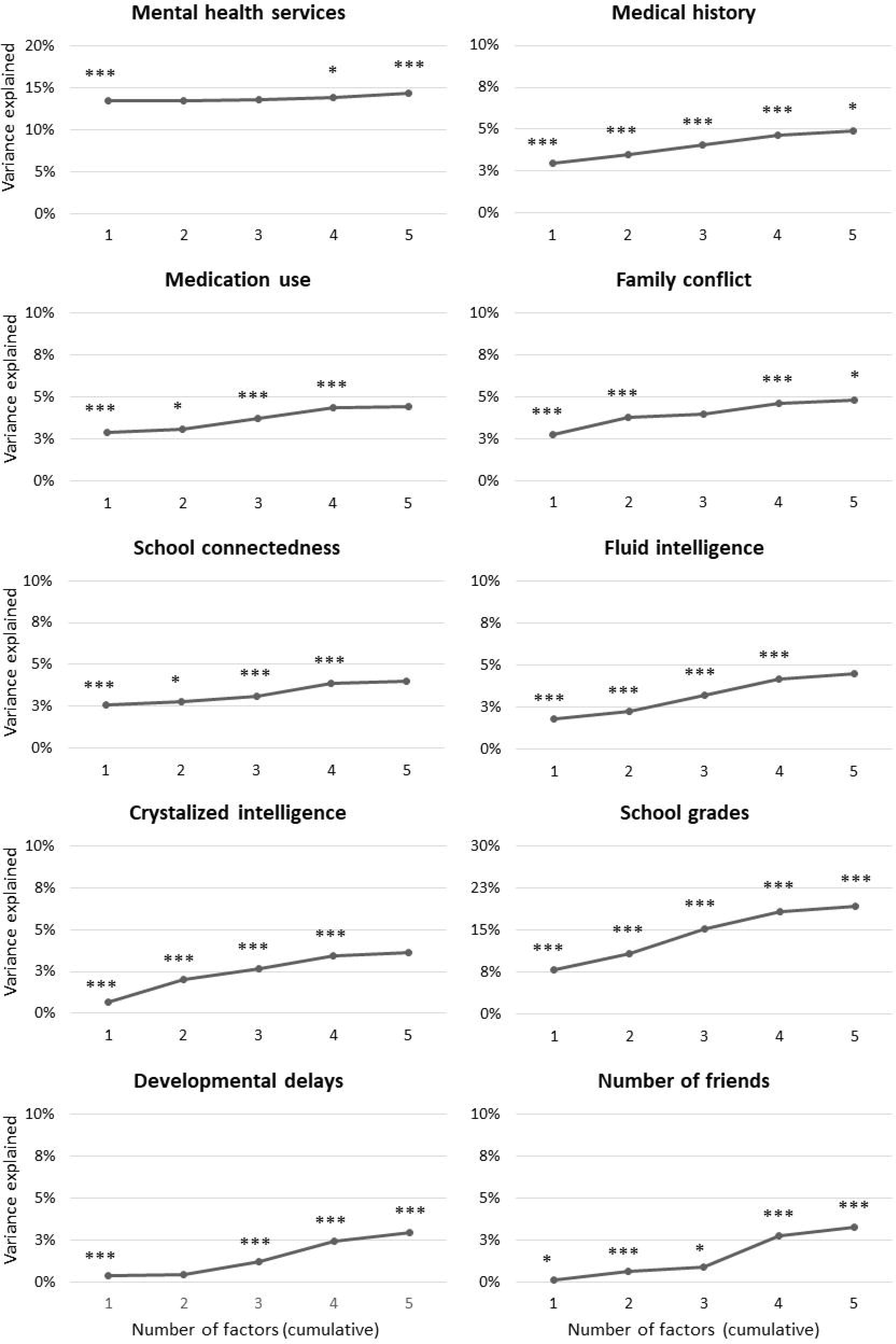
Cumulative explanatory power (R^2^) for a given factor structure (1- to 5-factor solutions) derived from CBCL data to predict validators.

Notes: Asterisks indicate significant change in R^2^ for that structure versus all simpler structures combined (\**P*<.05, \*\**P*<.01, \*\*\**P*<.001). CBCL=Childhood Behavior Checklist.

In the 5-factor solution, the highest correlation for family conflict was with the externalizing factor (*r*=.20) (Supplementary Table 3). Crystalized intelligence, fluid intelligence, and average grades showed the highest correlations with the neurodevelopmental factor (*r* between -.12 and -.37). Number of friends and developmental delays were most associated with detachment (*r*=−.15 and .12). In contrast, mental health services, medication use, medical history, and school connectedness showed generally non-specific correlations with psychopathology dimensions.

## 4. DISCUSSION

This study provides the most comprehensive examination of the hierarchy of psychopathology spectra to-date, in analyzing a wide range of symptoms and maladaptive behaviors, systematically explicating it across multiple hierarchical levels, considering both children and adults, and validating the structure against various clinically-relevant measures. In children, we found five spectra at the lowest level of the hierarchy: internalizing, somatoform, detachment, externalizing, and neurodevelopmental. In adults, we observed the same dimensions, except for separate disinhibited and antagonistic factors instead of a single externalizing factor at the lowest level. We further found substantial familiality of the identified psychopathology factors, largely explained by familial aggregation of the p-factor. Yet, the five childhood dimensions also showed specific links to corresponding parental dimensions. The p-factor was sufficient to account for some clinical validators (e.g., service utilization), but all five dimensions were needed to explain other validators, such as developmental delays, and social and educational functioning. These findings support the value of explicating multiple higher-order dimensions of psychopathology. They further suggest that the neurodevelopmental spectrum should be considered for inclusion in dimensional models of both childhood and adult psychopathology. Overall, the identified hierarchy depicts robust and informative dimensional phenotypes for the ABCD study baseline, paving the way for future research on this cohort.

In both children and adults, we observed that the p-factor at the top of the hierarchy separates into broad internalizing and broad externalizing spectra. These dimensions mirror the higher-order dimensions first identified by Achenbach and colleagues.^51^ Differentiation of the broad internalizing factor at lower hierarchical levels produced the emergence of narrower internalizing, detachment, and somatoform factors in both adults and children. These dimensions are consistent with prior studies identifying them among major dimensions of psychopathology.^8, 25, 51, 52^ From the broader externalizing factor, narrower externalizing and neurodevelopmental factors emerged in children, reflecting a separation of aggressive and rule-breaking behaviors from symptoms of inattention, hyperactivity, and related problems (e.g. clumsiness, speech problems, obsessions). In adults, three factors emerged from the broad externalizing spectrum: disinhibition, antagonism and neurodevelopmental. The additional dimension in adults is consistent with the hypothesis that psychopathology becomes more differentiated with age,^14, 33^ and with research showing a separation between disinhibited and antagonistic dimensions.^8, 53, 54^

Our findings are consistent with the HiTOP model by identifying internalizing, disinhibited, antagonistic, somatoform, and detachment dimensions. A thought disorder spectrum was not found, likely because of the very low prevalence of psychosis symptoms in this population-based sample. Conversely, we observed a neurodevelopmental dimension that includes inattention, hyperactivity, and clumsiness in adults, as well as autistic-like traits and atypical ideation (e.g. obsessions) in children. This dimension has previously been proposed as a neurodevelopmental spectrum^26^ and is consistent with initial factor analytic evidence in children.^27, 28, 35^ The emergence of this factor in adults is novel, as previous structural studies of adults have not considered enough neurodevelopmental problems to allow delineation of this dimension. This finding provides the strongest evidence to date for the inclusion of the neurodevelopmental spectrum in dimensional models of psychopathology.

By mapping multiple hierarchical levels, we show that the familial aggregation of psychopathological dimensions in parents and children on is largely accounted for by familial influences on the p-factor. This is consistent with the established pleiotropy in the genetic vulnerability to psychopathology^19, 55, 56^ and prior evidence of substantial heritability of the p-factor.^5^ Child p-factor also accounted for the majority of psychopathology-related variance in several validators, especially utilization of mental health services, which underscores the value of this general dimension for public health and planning of clinical services. However, more specific dimensions also proved to be informative. Familial aggregation between specific dimensions remained significant, albeit reduced, when controlling for p-factors, and all levels of the hierarchy showed incremental validity, with five dimensions necessary to maximize explanatory power of psychopathology for most criteria. This supports the importance of examining multiple levels of the psychopathology hierarchy, and is consistent with the views that fine-grained understanding of psychopathology is necessary to fully explicate its etiology^57, 58^ and identify maximally effective treatment.^59^ Further, different dimensions were most important for different validators. For example, the neurodevelopmental dimension had particularly strong links to intelligence and academic achievement, consistent with previous evidence,^60, 61^ and the externalizing factor with family conflict, as expected.^40^ These results confirm previous studies showing that both a general factor and specific dimensions are necessary for characterizing youth psychopathology,^19^ school grades, school and neighborhood deprivation,^14^ and executive functioning.^27^ They are inconsistent with studies linking cognitive abilities primarily to the p-factor,^11, 41^ potentially because these studies did not model the neurodevelopmental dimension, the strongest correlate of fluid and crystalized intelligence in this study.

The present study had the following limitations. First, it was limited to one assessment system, thus generalizability of the findings needs to be tested with other measures. Nevertheless, the hierarchy is largely consistent with previous studies using different measures^21, 23, 30, 53^ suggesting at least partial generalizability. Second, only one parent completed both the CBCL about the child and the ASR about themselves, which may have inflated the similarity between childhood and adult psychopathology structures due to rater biases. Although this limitation is common to much of the existing literature on parent and offspring psychopathology when children are too young to provide comprehensive self-reports, and a number of our validators were objective (e.g. cognitive testing) or self-report (e.g. number of friends) measures, future research should replicate the current results with child self- and additional co-informant reports. Third, only one time point was included, as longitudinal data were not yet available from the ABCD study at the time of writing. Future waves of data in this unique sample will provide the unprecedented opportunity to examine the hierarchy of psychopathology over the course of development and the predictive validity of childhood factors on a variety of adolescent and young adult outcomes.

In conclusion, the present results clarify the hierarchy of psychopathology dimensions in children and adults using data from one of the largest initiatives to study youth development and psychopathology to date. The study replicates higher-order dimensions identified previously,^8^ and suggests addition of the neurodevelopmental spectrum to dimensional models of psychopathology. The identified higher-order dimensions represent valid constructs able to explain various clinically-relevant risk factors and outcomes, such as developmental delays and academic achievement. Our investigation further provides a guide for future research to use these higher-order psychopathology dimensions in the ABCD sample. New data releases will allow researchers to replicate current results and apply the identified hierarchy to additional clinical, functional, and neuroimaging measures to study the interplay of psychopathological dimensions with adolescent development.

## ACKNOWLEDGMENTS

Drs Michelini and Kotov are funded by National Institute of Mental Health (NIMH) award number MH117116. Data used in the preparation of this article were obtained from the Adolescent Brain Cognitive Development (ABCD) Study (https://abcdstudy.org), held in the NIMH Data Archive (NDA). The ABCD Study is supported by the National Institutes of Health (NIH) and additional federal partners under award numbers U01DA041022, U01DA041025, U01DA041028, U01DA041048, U01DA041089, U01DA041093, U01DA041106, U01DA041117, U01DA041120, U01DA041134, U01DA041148, U01DA041156, U01DA041174, U24DA041123, and U24DA041147. A full list of federal partners is available at https://abcdstudy.org/federal-partners.html. A listing of participating sites and a complete listing of the study investigators can be found at https://abcdstudy.org/principal-investigators.html. This manuscript reflects the views of the authors and may not reflect the opinions or views of the NIH or ABCD consortium investigators. ABCD consortium investigators designed and implemented the study and/or provided data but did not necessarily participate in analysis or writing of this report. The authors would like to thank Avshalom Caspi, PhD (Duke University; King’s College London) and Terrie Moffitt, PhD (Duke University; King’s College London) for their helpful comments on an earlier draft of this manuscript.

