## Supplementary Method 1 for "Delineating and validating higher-order dimensions of psychopathology in the Adolescent Brain Cognitive Development (ABCD) study"

**Further details on the measures**

1. ***Childhood psychopathology***

Emotional and behavioral problems in the children were assessed with the Child Behavior Checklist (CBCL) (1), completed by parents during in-person assessments. The CBCL includes 112 problem items rated on the scale 0 = Not True, 1 = Somewhat or Sometimes True, and 2 = Very True or Often True. From these items it is possible to derive two higher-order internalizing and externalizing total scores (2), as well as eight syndrome scales: anxious/depressed, withdrawn/depressed, somatic complaints, social problems, thought problems, attention problems, rule-breaking behavior, and aggressive behavior (1). We analyzed individual items rather than pre-defined scales.

2. ***Adult psychopathology***

Emotional and behavioral dimensions relevant to adult psychopathology were measured in the primary caretaker parent with the Adult Self Report (ASR) (3). It includes 120 problem items rated on the same scale as the CBCL, between 0 and 2. Similar to the CBCL, the ASR items can be aggregated into two higher-order internalizing and externalizing total scores, as well as eight syndrome scales: anxious/depressed, withdrawn, somatic complaints, thought problems, attention problems, rule-breaking behavior, aggressive behavior, and intrusive (3). We analyzed individual items.

3. ***Developmental History***

To measure developmental delays in motor and speech acquisition, we created a composite score of two questionnaire items completed by parents asking about motor and speech development compared to most children, on the following scale: 1 = Much earlier; 2 = Somewhat earlier; 3 = About average; 4 = Somewhat later; 5 = Much later (4).

4. ***Peer and family environment***
   4.1. Number of friends: the child’s peer relationships were assessed with a total score from a 4-item questionnaire asking the child about the number of friends he/she has (5). Two item on the number of close female and male friends were rescored with the following windows: 0 = 0; 1 = 1; ... 10 = 10; 11 to 100 = 11. Two items on the number of other female and male friends were rescored with the following windows: 0 = 0; 1 = 1; ... 10 = 10; 11 to 15 = 11; 16 to 20 = 12; 21 to 25 = 13; 26 to 30 = 14; 31 to 100 = 15. The total of the 4 items was coded as a count variable.
   4.2. Family conflict: parent-reported conflict within the family was assessed with the total score of 9 items from the Family Conﬂict subscale of the Moos Family Environment Scale (FES) (5, 6).
5. ***School environment and grades***
   5.1. School connectedness: the child’s connectedness to his/her school in relation to teachers, classroom environment, personal involvement in school, and academic goals was obtained by creating a composite score from 12 items of the School Risk and Protective Factors Survey from the PhenX Toolkit, completed by the child (5).
   5.2. School grades: parents reported on what grades the child gets on average, on the following scale: 6 = A's / Excellent; 5 = B's / Good; 4 = C's / Average; 3 = D's / Below Average; 2 = F's / Struggling a lot; 1 = ungraded.
6. ***Service utilization***
   6.1. Medical history: parents were asked two questions from the MAGIC Health Services Utilization Questionnaire (7) asking whether the child has seen any health professional (other than regular check-ups) in the past year and in their lifetime (4). A sum of this two items rated on a dichotomous scale (1 = Yes, 0 = No) was used in analysis.
   6.2. Mental health services: parents were asked whether the child ever received mental health or substance abuse services (4).
   6.3. Medication history: parents were asked whether their child took any type of medication in the two weeks prior to the visit (4).
7. ***Cognitive abilities***

The National Institute of Health (NIH) Toolbox was completed by each child to obtain measures of cognitive abilities. To assess performance and verbal cognitive abilities in the present study we used, respectively, the NIH Toolbox Crystalized Intelligence Composite (sum of scores from Picture Vocabulary Test and Oral Reading Recognition Test) and a Fluid Intelligence Composite (sum of scores from List Sorting Working Memory Test, Pattern Comparison Processing Speed Test, Picture Sequence Memory Test, Flanker Task, and Dimensional Change Card Sort Test) (8). These composite scores show good test-retest reliabilities and validity in children (8, 9).

**Preliminary analyses of CBCL and ASR items**

To address problems in EFA with items that were not analyzable due being endorsed too infrequently or being too-highly correlated with other items, we took two approaches. Firstly, we examined frequencies of all items included in CBCL and ASR, and removed items which frequency was too low (<1% rated as 1 or 2). Secondly, to address high inter-item correlations, which can distort factor structure, we aggregated items that were highly correlated (polychoric r>.75) into composites by averaging scores and then rounding to the nearest integer, thus preserving the trichotomous rating.

The following CBCL items were removed because of low frequency: “Drinks alcohol without parents' approval”, “Problems with eyes (not if corrected by glasses)”, “Other (physical problems without known physical cause)”, “Sets fire”, “Sexual problems”, “Smokes, chews, or sniffs tobacco”, “Truancy, skips school”, “Uses drugs for non-medical purposes (don't include alcohol or tobacco)”, “Wishes to be of opposite sex”. The following composites were created: Attacks/threatens (“Physically attacks people”, “Threatens people”); Destroys (“Destroys his/her own things”, “Destroys things belonging to his/her family or others”, “Vandalism”); Disobeys rules (“Disobedient at home”, “Disobedient at school”, “Breaks rules at home, school or elsewhere”); Steals (“Steals at home”, “Steals outside the home”); Peer problems (“Doesn't get along with other kids”, “Not liked by other kids”); Distracted (“Can't concentrate, can't pay attention for long”, “Inattentive or easily distracted”); Hallucinations (“Hears sound or voices that aren't there”, “Sees things that aren't there”); Sex play (“Plays with own sex parts in public”, “Plays with own sex parts too much”); Weight problems (“Overeating”, “Overweight”). EFAs were run on the remaining 90 original and 9 composite items.

The following ASR items were removed because of low frequency: “I steal”, “I threaten to hurt people”. The following composites were created: Anxious (“I am nervous or tense”, “I am too fearful or anxious”); Overt aggression (“I damage or destroy my things”, “I damage or destroy things belonging to others”, “I physically attack people”); Money management (“I have trouble managing my money or credit card”, “I fail to pay my debts or meet other financial responsibilities”); Nausea (“Nausea, feel sick”, “Vomiting, throwing up”); Suicidality (“I deliberately try to hurt or kill myself”, “I think about killing myself”); Mood swings (“My moods swing between elation and depression”, “My moods or feeling change suddenly”); Hallucinations (“I hear sounds and voices that other people think aren't there”, “I see things that other people think aren't there”); Oddness (“I do things that other people think are strange”, “I have thoughts that other people would think are strange”). EFAs were run on the remaining 101 original and 8 composite items.
