## Supplementary figures and images for "Delineating and validating higher-order dimensions of psychopathology in the Adolescent Brain Cognitive Development (ABCD) study"

### Supplementary Figure 1

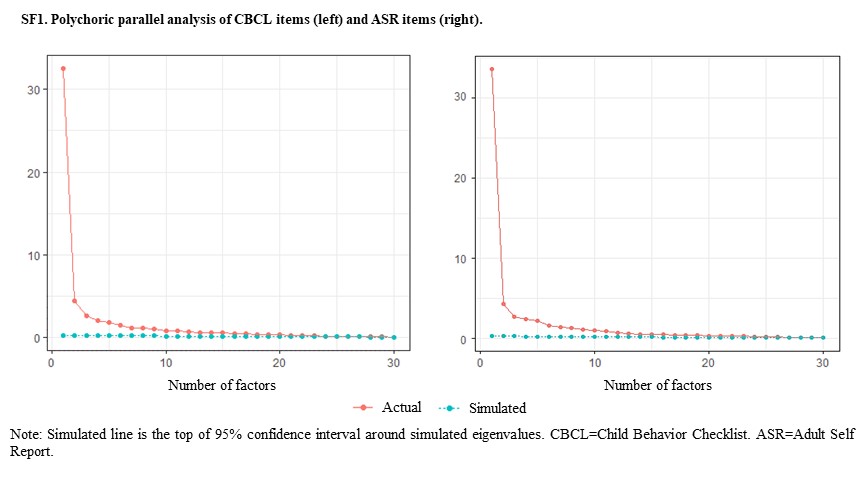
